# Hibernation slows epigenetic aging in yellow-bellied marmots

**DOI:** 10.1101/2021.03.07.434299

**Authors:** Gabriela M. Pinho, Julien G. A. Martin, Colin Farrell, Amin Haghani, Joseph A. Zoller, Joshua Zhang, Sagi Snir, Matteo Pellegrini, Robert K. Wayne, Daniel T. Blumstein, Steve Horvath

**Affiliations:** Department of Ecology and Evolutionary Biology, University of California, 621 Young Drive South, Los Angeles, CA 90095–1606, USA; Department of Biology, University of Ottawa, 30 Marie Curie, Ottawa, ON K1N 6N5, Canada; Department of Molecular, Cell and Developmental Biology, University of California, Los Angeles, CA, USA; Human Genetics, David Geffen School of Medicine, University of California, Los Angeles CA 90095, USA; Department of Evolutionary & Environmental Biology, Institute of Evolution, University of Haifa, Haifa, Israel; Rocky Mountain Biological Laboratory, Box 519, Crested Butte, CO 81224, USA; Biostatistics, Fielding School of Public Health, University of California, Los Angeles, Los Angeles, California, USA

**Keywords:** DNA methylation, epigenetic clock, epigenetic pacemaker, seasonality, torpor

## Abstract

Species that hibernate live longer than would be expected based solely on their body size. Hibernation is characterized by long periods of metabolic suppression (torpor) interspersed by short periods of increased metabolism (arousal). The torpor-arousal cycles occur multiple times during hibernation, and it has been suggested that processes controlling the transition between torpor and arousal states cause aging suppression. Metabolic rate is also a known correlate of longevity, we thus proposed the ‘hibernation-aging hypothesis’ whereby aging is suspended during hibernation. We tested this hypothesis in a well-studied population of yellow-bellied marmots (*Marmota flaviventer*), which spend 7-8 months per year hibernating. We used two approaches to estimate epigenetic age: the epigenetic clock and the epigenetic pacemaker. Variation in epigenetic age of 149 samples collected throughout the life of 73 females were modeled using generalized additive mixed models (GAMM), where season (cyclic cubic spline) and chronological age (cubic spline) were fixed effects. As expected, the GAMM using epigenetic ages calculated from the epigenetic pacemaker was better able to detect nonlinear patterns in epigenetic age change over time. We observed a logarithmic curve of epigenetic age with time, where the epigenetic age increased at a higher rate until females reached sexual maturity (2-years old). With respect to circannual patterns, the epigenetic age increased during the summer and essentially stalled during the winter. Our enrichment analysis of age-related CpG sites revealed pathways related to development and cell differentiation, while the season-related CpGs enriched pathways related to central carbon metabolism, immune system, and circadian clock. Taken together, our results are consistent with the hibernation-aging hypothesis and may explain the enhanced longevity in hibernators.

## Introduction

Aging is a poorly understood natural phenomenon, characterized by an age-progressive decline in intrinsic physiological function^1,2^. The high variation in disease and functional impairment risk among same-age individuals shows that biological age is uncoupled from chronological age^3–5^. Some individuals age at faster rates than others, and little is known about the causes of this inter-individual variance in biological aging rates^6,7^. To this end, researchers have been attempting to develop biomarkers of aging^4,8^. DNA methylation (DNAm) based age estimators, also known as epigenetic clocks (ECs), are arguably the most accurate molecular estimators of age^3,9–12^. An EC is usually defined as a penalized regression, where chronological age is regressed on methylation levels of individual cytosines^13^. The EC has been successfully used to study human aging, and is becoming increasingly used to study aging in other species^14–20^.

Biological processes underpinning ECs remain to be characterized^11^. Age-adjusted estimates of epigenetic age (epigenetic age acceleration) are associated with a host of age related conditions and stress factors, such as cumulative lifetime stress^21^, smoking habits^22,23^, all-cause mortality^24–28^, and age-related conditions/diseases^13,26,29,30^. These associations suggest that epigenetic age is an indicator of biological age^5,31^. In fact, measures of epigenetic aging rates are associated with longevity at the individual level as well as across mammalian species^6,17^. Several studies present evidence that long-lived species age more slowly at an epigenetic level^19,20,31–33^. The link between the epigenetic aging and biological aging is further reinforced by the observation that treatments known to increase lifespan significantly slow the EC^15,17^.

Longevity is related to body size, but some species have longer lifespans than expected based on their body size^34,35^. A characteristic of long-lived species is the ability to engage in bouts of torpor^36,37^. Torpor is a hypometabolic state characterized by a dramatic decrease in gene transcription and translation rates^38–43^. During hibernation, torpor bouts are interspersed by short periods of euthermy (< 24 h), when gene expression occurs and metabolism is fully recovered^41,44^. Some of the physiological stresses from the cyclic transition between deep torpor and euthermy are similar to the ones experienced by the aging body (e.g., oxidative stress), and promote responses in cellular signaling pathways that are essential for both longevity and torpor survival^36,45^. Thus, the cellular and molecular stress responses associated with torpor-arousal cycles and long periods of inactivity may suppress aging^36,45^.

Hibernation is mostly comprised of long periods of metabolic suppression, and overall metabolic rate reduction is associated with longevity^45–47^. Therefore, we hypothesize that aging is reduced during hibernation which we refer to as the hibernation-aging hypothesis. Specifically, a species that engages in torpor may periodically “suspend” aging, as previously suggested^36^. With this rationale, we predict that the epigenetic aging is faster during the active season and slower during hibernation. We test this prediction in yellow-bellied marmots (*Marmota flaviventer*), which spend 7-8 months per year hibernating^48^. Torpor bouts represent 88.6% of the yellow-bellied marmot hibernation period, resulting in an average energy saving of 83.3% when compared to the energetic expenditure of an euthermic adult^48,49^.

## Methods

All samples were collected as part of a long-term study of a free-living population of yellow-bellied marmots in the Gunnison National Forest, Colorado (USA), where marmots were captured and blood samples collected biweekly during the their active season (May to August^50^). Data and samples were collected under the UCLA Institutional Animal Care and Use protocol (2001-191-01, renewed annually) and with permission from the Colorado Parks and Wildlife (TR917, renewed annually).

Individuals were monitored throughout their lives, and chronological age was calculated based on the date at which juveniles first emerged from their natal burrows. We only used female samples because precise age for most adult males is unavailable since males are typically immigrants born elsewhere^51,52^. We selected 160 whole blood samples from 78 females with varying ages. From these, DNA methylation (DNAm) profiling worked well for 149 samples from 73 females with ages varying from 0.01 to 12.04 years.

Genomic DNA was extracted with Qiagen DNeasy blood and tissue kit and quantified with Qubit. DNAm profiling was performed with the custom Illumina chip HorvathMammalMethylChip40^53^. This array, referred to as mammalian methylation array, profiles 36 thousand CpG sites in conserved genomic regions across mammals. From all probes, 31,388 mapped uniquely to CpG sites (and its respective flanking regions) in the yellow-bellied marmot assembly (GenBank assembly accession: GCA_003676075.2). We used the SeSaMe normalization method to estimate β values for each CpG site^54^.

Two model approaches were used to study epigenetic aging in marmots: the epigenetic clock^9,10^ and the epigenetic pacemaker^55–58^. Both models are described below.

### Epigenetic clock (EC)

Under the EC a linear correlation with age is determined by attempting to fit a single coefficient to each CpG site. We fitted a generalized linear model with elastic-net penalization^59^ to the chronological-age and β-value data sets using the glmnet v.4.0-2 package in R^60^. Alpha was set to 0.5, which assigns ridge and lasso penalties with same weight. The elastic-net penalization limits the impact of collinearity and shrinks irrelevant coefficients to zero. This method estimates coefficients that minimize the mean squared error between chronological and predicted ages, and performs an automatic selection of CpG sites for age prediction. We applied a 10-fold cross validation to select the model with lowest error based on the training set. Predicted ages were scored for samples not included in the training set of the model (code will be available in supplementary material). In this respect, the predicted age was estimated for groups of ~14 samples, resulting in 11 EC models. These models comprised 360 sites, and the average coefficient per site and intercept will be available in the supplementary material.

### Epigenetic pacemaker

While ECs are used to estimate the age of a sample based on weighted sums of methylation values, the epigenetic pacemaker (EPM) models the dynamics of methylation across the genome. To accomplish this, it models each individual CpG site as a linear function of an underlying epigenetic state of an individual. This epigenetic state changes with time in a nonlinear fashion, and we are therefore able to use this paradigm to identify periods with variable rates of methylation changes throughout lifespan. The EPM assumes that the relative increase/decrease rate of methylation levels among sites remains constant, but the absolute rates can be modified when rates at all sites change in synchrony^56–58^. The optimum values of methylation change rate and initial methylation level per site, as well as the epigenetic state per sample, are calculated through iterations implemented in a fast conditional expectation maximization algorithm^61^ to minimize the residual sum of squares error between known and estimated methylation levels (β values). Thus, the epigenetic state is an estimate of age that, given the methylation rates and initial methylation levels for each site, minimizes the differences between known and estimated methylation levels in a specific sample for all sites included in the model. We selected sites to use in the EPM based on the absolute Pearson correlation coefficients (*r*) between chronological age and methylation levels per site^57,58^. All sites with *r* > 0.7 were included, which resulted in 309 sites. A 10-fold cross validation was used to estimate epigenetic states (supplementary material). We report the rate and intercept values per site from the EPM using all data as training set (no cross validation, supplementary material).

### Hibernation-aging hypothesis

We fitted two Generalized Additive Mixed Models (GAMM) with the EPM- or the EC-estimated epigenetic age as dependent variable. For both GAMMs, fixed effects included a cubic spline function for chronological age, and a cyclic cubic spline function for day of year. We also tested for the interaction between these two variables (using tensor product interaction with a cubic spline). Individual identity was added as random effect. Day of year varied from 1 to 365, with 1 representing 1 May and 365 representing 30 April.

We used simulations to estimate the type 1 error and the potential power to detect a hibernation-aging effect given the limitations of our sample collection. Specifically, blood samples could only be collected during the active season, instead of throughout the year. Our earliest sample was collected on 27 April and the latest on 20 August. We simulated two different traits: (1) a trait that increases linearly with age independently of the season; and (2) a trait that increases during the summer but not during the winter. The daily rate of increase for the first trait was set at 0.004, to simulate data with a similar range to the observed EPM data. For the second trait, the rate of increase was set to zero during winter (16 Sept to 17 April, days 139-352 using 1 May as reference). The simulation assumed that the active season was 150 days long starting on 18 April (day 353) and finishing on 15 Sept (day 138). The rate of increase during the active season was set as 0.0164 (0.004 / 365 * 150) so that the annual rate of increase was similar between the two simulated traits. Our simulation was parameterized using among-individual and residual variance from the EPM. We performed these simulations using field data (day of sample collection, age in days, birth date, and number of samples), and estimated the significance of the seasonal effect (cyclic spline with days since 1 May). We repeated this procedure 1000 times for both traits. The proportion of simulations on trait 1 (no seasonal effect) that were significant indicated our type 1 error. The proportion of simulations on trait 2 (seasonal effect) that were significant was an indication of the power to detect this effect.

We evaluated GAMMs by checking convergence, concurvity between fixed effects and the autocorrelation of deviance residuals. We also checked model fit by plotting fitted with observed epigenetic state and visually inspected qq plots and histograms of deviance residuals, plots of deviance residuals with fitted values, and plots of deviance residuals with explanatory variables. GAMMs were fitted and checked using the mgcv R package v. 1.8^62^. All analysis and figures were developed in R v.3.6.3^63^ in RStudio v. 1.2.5033^64^, python v.3.7.4^65^, Jupyter notebook v.6.0.3^66^, ggplot2 v.3.3.0.9^67^, and ggpubr v.0.2.5^68^.

### Influence of chronological age and seasons on methylation levels per CpG site

We performed additional analyses to identify which CpG sites were associated with age and seasonality. We fitted a Generalized Additive Model (GAM) per CpG site, where methylation level was the dependent variable. The independent variables were a cubic spline function for chronological age and a cyclic cubic spline function for day of year.

Since epigenome wide association studies of age (EWAS) have been more commonly used to identify CpG sites related to chronological age, we performed a linear regression per CpG site. Each model had methylation level as dependent variable and chronological age as independent variable. Significance thresholds were set to 1×10^-5^.

### CpG site enrichment analysis

Gene enrichment was performed with the Genomic Regions Enrichment Tool v.3.0.0 (GREAT hypergeometric test^69^). GREAT analyzes the potential cis-regulatory role of the non-coding regions with CpG sites of interest, and identifies which pathways are overrepresented in the data. To associate CpGs with genes, we used the “Basal plus extension” association with a maximum window distance between the CpG and the genes of 50 kb. GREAT tests the observed distribution of CpG neighboring genes against the expected number of sites associated with each pathway due to their representation in the mammalian array (background set). Since GREAT requires a high quality annotation, we used the respective locations of the marmot sites on the human assembly (GRCh37), and therefore only used sites mapped to conserved genes between marmots and humans. Two data sets were analyzed: sites associated with chronological age and with day of year. The alignment and annotation methods are described in the mammalian methylation array method paper^53^.

## Results

The epigenetic aging models developed with the epigenetic clock (EC) and the epigenetic pacemaker (EPM) were both highly accurate (Figure 1), showing high correlations between epigenetic and chronological age (*r* = 0.98 and 0.92, respectively). The EC used 360 CpG sites, whereas the EPM used 309 sites. There was an overlap of 72 sites between the two models.

**Figure 1:**
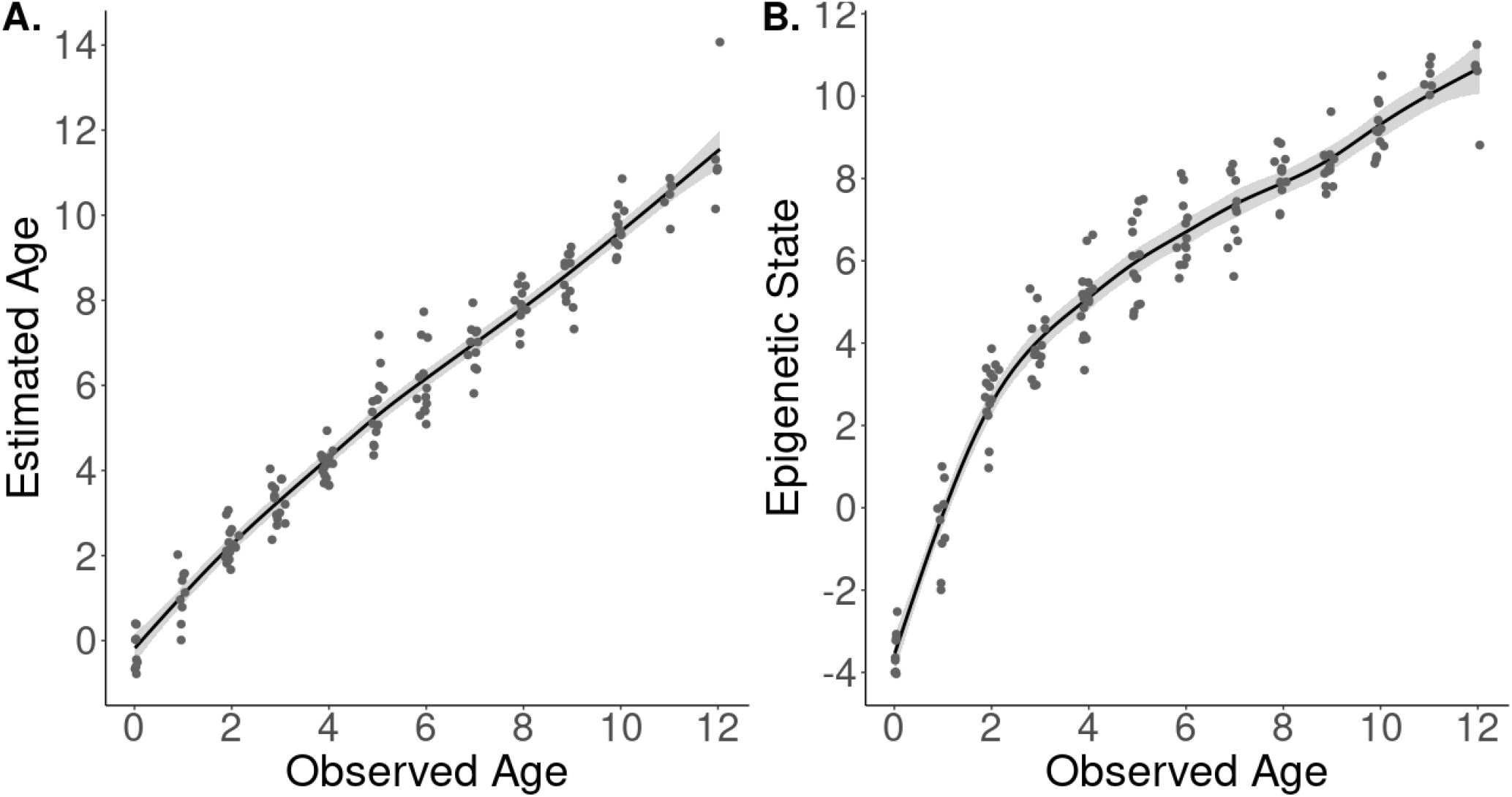
Epigenetic aging models for a wild population of yellow-bellied marmots developed from the epigenetic clock (A), and the epigenetic pacemaker (B). Points represent samples from individuals of known age at the sampling moment (Observed Age), and y-axis represent the epigenetic age calculated by each model. Trend lines were developed by fitting cubic splines.

The GAMM fitted to the predicted age from the EC explained 96.6% of the variation (Adj. R^2^) and had a residual variance of 0.346. The random effect of individual identity had an intercept variance of 0.009. The age spline was significant (F = 1225.76, p < 0.0001) and the cyclic spline for days since 1 May was not significant (p = 0.78). The tensor interaction smooths was also not significant (p = 0.11). Details of this model will be described in the supplementary material.

The GAMM fitted to the epigenetic state data explained 95.6% of the variation and had a residual variance of 0.284. The random effect of individual identity had an intercept variance of 0.332. Both smooth terms significantly influenced marmot epigenetic state (p < 0.005, Table 1), but the interaction between them was not significant (p = 0.44). The effect of chronological age and day of year result in a particular pattern of epigenetic state change (Figure 2A). The partial effect of day on epigenetic state shows an increase in epigenetic state during the summer and suggests a reversal of such changes during the winter (Figure 2B). Moreover, the rate of epigenetic state increase is the highest in the mid-point of the active season. The partial effect of chronological age shows that the epigenetic state increases at a higher rate until females reach 2-years old, followed by a deceleration as individuals become older (Figure 2C).

**Figure 2.**
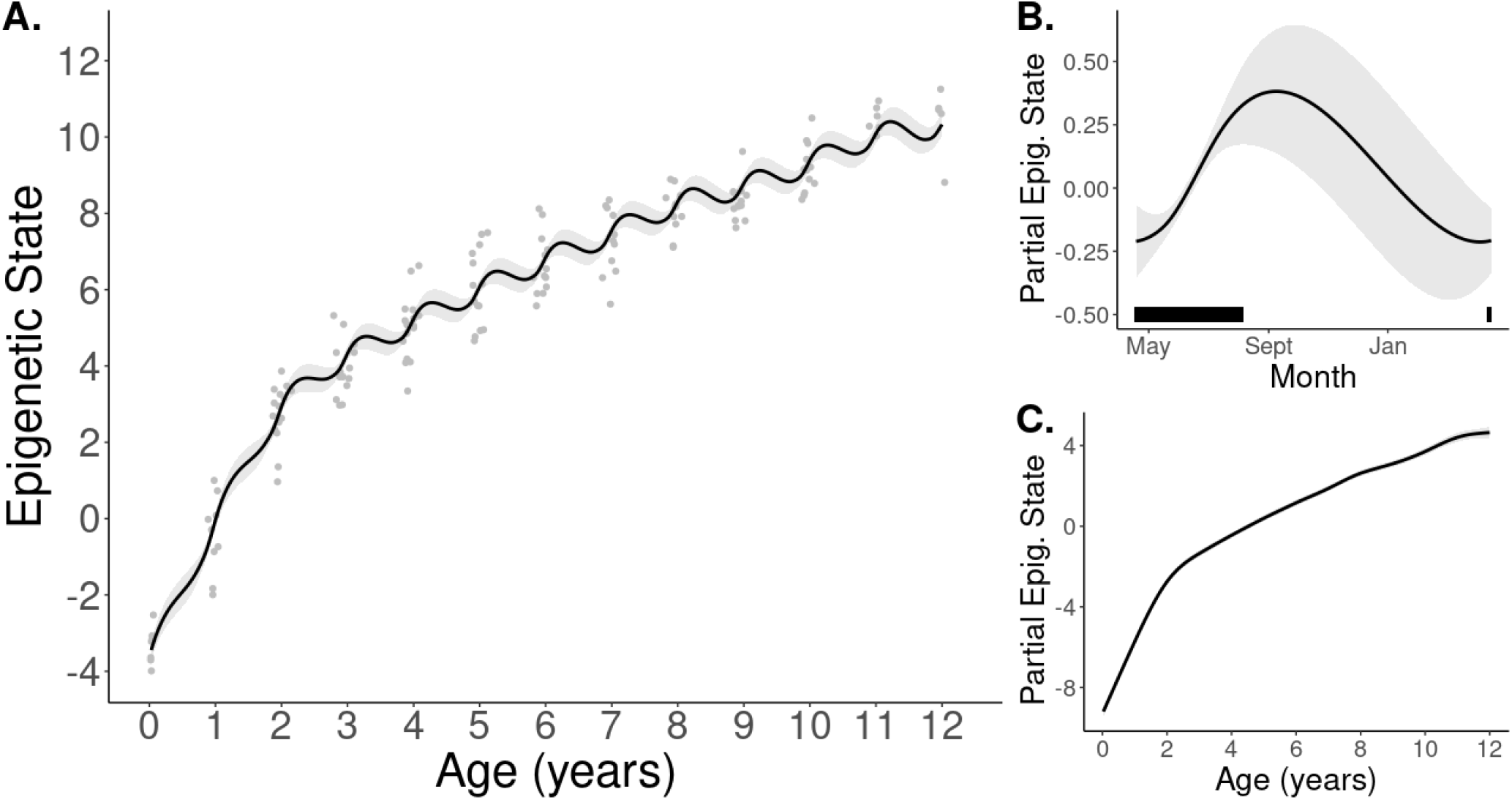
Visualization of the generalized additive mixed model with epigenetic states generated from the epigenetic pacemaker model using CpG sites highly correlated to chronological age (absolute *r* > 0.7). A) Changes in the epigenetic state (or epigenetic age) as individuals age. Points are actual data, while lines are the predictions from the model. B) Predictions generated with the partial effect of date of year (cyclic cubic smoother spline) on epigenetic state. The black horizontal bar represents when samples were collected and most of the marmot active season. C) Predictions generated with the partial effect of chronological age (cubic smoother spline) on epigenetic state. Buffers illustrate the 95% confidence intervals.

**Table 1.** Output from the generalized additive mixed model using epigenetic states (or epigenetic ages) as dependent variable. Epigenetic states were estimated from epigenetic pacemaker models (EPM). Age: individual chronological age in years calculated from the first time an individual emerged from their mother’s burrow to the date they were captured. Date: day of the year (values varied from 1 to 365, with 1 representing 1 May and 365 representing 30 April).

|  | **Estimate** | **Std. Error** | **t value** | **p-value** |
| --- | --- | --- | --- | --- |
| **Intercept** | 5.53 | 0.09 | 63.39 | <0.0001 |
| **Smooth terms:** |  |  |  |  |
|  | **edf** | **Ref.df** | **F** | **p-value** |
| **Age**<br>**(cubic spline)** | 8.43 | 8.43 | 291.82 | <0.0001 |
| **Date**<br>**(cyclic spline)** | 1.39 | 8.00 | 1.08 | 0.003 |
| **Age**<br>**(cubic spline)**<br>**×**<br>**Date**<br>**(cyclic spline)** | 0.00 | 12.00 | 0.00 | 0.440 |
| **Random effect:** |  |  |  |  |
|  | **Intercept** | **SD Residual** |  |  |
| **Animal ID** | 0.576 | 0.532 |  |  |

### Simulation

From the 1000 GAMMs fitted to data simulated with a seasonal effect, 76.5% found a significant effect of seasons, indicating high power to detect a seasonal effect given the simulated parameters and our data structure. From the 1000 GAMMs fitted to data simulated with a constant linear age effect, 7.3% had a significant season effect, indicating a slightly higher type 1 error than expected (5%). Based on this result from the simulations with no seasonal effect, we calculated a new critical value for the probability that respects the 5% type 1 error rate by estimating the 0.05 quantile of the p-value distribution from a null model. The 0.05 quantile was 0.0344, which can be taken as the critical value with which to estimate the significance of a seasonal effect. The p-value for seasonal effects on our data is < 0.0344 and therefore is considered significant. From this, we concluded that our results were not driven by our sampling.

### Age-related CpGs

In the EWAS of chronological age, the methylation level of 6,364 CpGs were significantly (p < 10^-5^) associated with chronological age. In the GAMs per site, the age effect (cubic smoother spline) was significant in 6,303 sites, which largely overlapped with EWAS of age (Figure 3D). From the 5,841 sites that overlapped between the EWAS and the age effect, 66% (3,827 sites) had effective degrees of freedom (edf) values larger than 2 for the age effect in the GAMs. The edf measures the complexity of the curve, and these results imply that most CpG sites have a non-linear relationship with chronological age. Top age-related CpGs in both EWAS and GAMs were located on NR2F1 and EVX2 downstream regions (Figure 3AB). The promoters of EN1 and HOXD10 were also hypermethylated with age. Age-related sites uniquely identified by GAMs were proximal to FAM172A intron (hypermethylated), and hypomethylated in both CSNK1D 3’UTR and HNRNPC intron.

**Figure 3.**
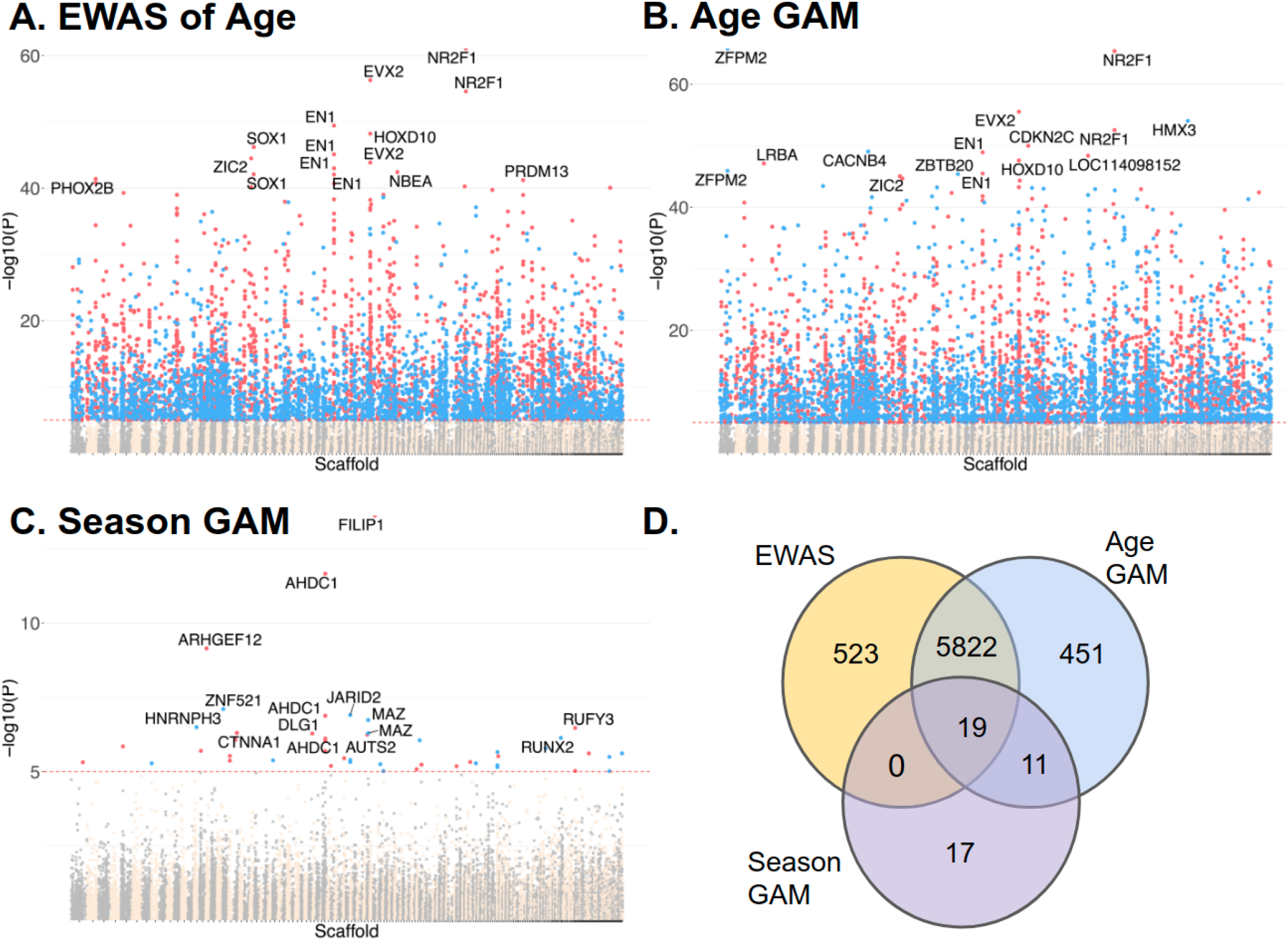
Associations of CpG sites with chronological age and seasons (day of the year) in blood of yellow bellied marmots (*Marmota flaviventer*). A, C) Manhattan plots visualizing log transformed p-values. The y-axis reports p values for two fixed effects of the Generalized Additive Models of individual cytosines (dependent variable): (A) chronological age (cubic spline function) and (C) day of year (cyclic cubic spline function). B) The y-axis reports p values for the epigenome-wide association (EWAS) of chronological age. The CpG sites coordinates were estimated based on the alignment of Mammalian array probes to yellow-bellied marmot genome assembly. The direction of associations with chronological age is highlighted for the significant sites (p < 10^-5^) with red for hypermethylated and blue for hypomethylated sites. Note that the season effect is cyclical, and we show the direction od association with chronological age for the active season. D) Venn diagram showing the overlap of significant CpG sites between EWAS and GAMs.

The 3,914 CpGs used in the enrichment analysis were located in both genic and intergenic regions relative to transcriptional start sites, with a higher proportion located at promoter regions than in the background (supplementary material). Most CpGs in promoter regions were hypermethylated with age (supplementary material). DNAm aging in marmots was proximal to polycomb repressor complex targets (PRC2, EED) with H3K27ME3 marks (supplementary material), which is a consistent observed pattern in all mammals^70^. The enriched pathways were largely associated with development, cell differentiation and homeostasis.

### Season-related CpGs

The seasonal effect in the GAMs per site, measured with a cyclic cubic spline function of day of the year, was significantly associated with methylation in 47 CpG sites proximal to 37 genes. Most of the season-related CpGs were also associated with age (Figure 3D). Some of the top season- and age-related CpGs are proximal to FILIP1 exon, ARHGEF12 intron, ZNF521 intron, JARID2 exon, and AHDC1 intron (Figure 3C). The top season sites with no association with age are proximal to AHDC1 intron, MAZ exon, CTNNA1 exon, AUTS2 intron, and EFNA5 exon (Figure 3C). The AHDC1 intron seems to be an interesting region for further exploration because it is proximal to sites solely affected by season, to sites only related with age, and those influenced by both. Mutations in AHDC1 are implicated in obstructive sleep apnea (PMID 31737670), so this gene may play a role in sleep processes, and potentially hibernation.

Since the seasonal effect size is smaller and more nonlinear than the age effect (Figure 2), our power to identify sufficient season CpGs for enrichment analysis was limited by our sample size. Thus, we performed a second enrichment analysis in a CpG site set selected using a less conserved false detection rate correction (Benjamini–Hochberg FDR^71^). With this method, 206 CpGs were significantly affected by season, and 126 were used in the enrichment analysis. Some interesting biological functions in this set included pyruvate metabolism (GO:0006090), transporters of monocarboxylates (GO:0008028, GO:0015355), leukocyte migration (GO:0050900), and the circadian clock system (GO:0032922, P00015, MP:0002562).

## Discussion

Acquiring chronological-age data from wildlife is a daunting task, but age data has fundamental applications to behavioral ecology, evolutionary biology, and animal conservation^7,72^. Epigenetic clocks (ECs) promise to inform age estimates in wild and non-model organisms^14,18,72^. This is the first study to present epigenetic aging models for marmots, a fascinating animal model to study hibernation. We applied a validated platform for measuring methylation levels (mammalian methylation array^53^) to a unique collection of tissues—blood samples from known age, free-living animals—to investigate how aging is affected by active-hibernation cycles.

The epigenetic pacemaker (EPM) results showed a rapid change in epigenetic age until marmots reached 2-years old, their age of sexual maturity^51,73^. After reaching adulthood, epigenetic age change was more linear and slower, which is similar to the pattern observed in humans older than 20 years^57^. The pattern observed in marmot epigenetic aging is consistent with the notion that methylation remodeling is associated with key physiological milestones^33^. A logarithmic relationship between methylation change rate and chronological age may be a shared trait in mammals, and such a relationship has been described for multiple human tissues^9,55,57^ and species, including dogs^33^, mice^15^, and yellow-bellied marmots.

With regard to active and hibernation seasons, the EC model was unable to capture seasonal effects because it uses a penalized regression to relate the dependent variable (chronological age) to cytosines. The EPM is better equipped to detect non-linear and potentially cyclic patterns because it estimates the epigenetic state by minimizing the error between estimated and measured methylation levels^57,58^, which allows for a non-linear relationship of methylation levels with chronological age. Since aging rate is not constant throughout an individual’s lifespan^74,75^, the EPM is possibly more influenced by factors associated with biological aging^57^. In fact, methylation levels in most CpG sites had a non-linear relationship with chronological age in our models per CpG site.

According to the model that used EPM-estimated epigenetic age, biological aging slows during hibernation. Specifically, the clear delay in epigenetic-state changes during hibernation supports our hibernation-aging hypothesis. Interestingly, this hypothesis does not seem to hold for individuals prior to sexual maturity. Even though we observed a non-significant interaction between chronological age and day of year, our model predictions indicated a weaker deceleration in aging during hibernation for individuals in their first and second years of life (Figure 2A). Compared to adults, young marmots spend less time torpid during hibernation, have higher daily mass loss in deep torpor^48^, and may immerge into hibernation weeks later^76–78^. Indeed, thermoregulatory support from adults increases overwinter survival of young alpine marmots^79–81^. Thus, a weaker effect of slowed aging during hibernation in younger animals may be explained by their later hibernation start date in addition to an overall higher metabolic rate during hibernation.

Some of the physiological stresses experienced by individuals during hibernation are similar to those observed with aging, and therefore the molecular and physiological responses required for an individual to successfully hibernate may prevent aging^36,45^. Additionally, hibernation combines conditions known to promote longevity^36,45,82^, such as food deprivation (calorie restriction^83–85^), low body temperature^82,86–88^, and reduced metabolic rates^45^. Conceivably, these factors may also be associated with the slower marmot aging observed in the beginning and end of their active season (Figure 2B). Marmots in early Spring and late Fall have limited calorie intake^78,89^, reduced overall activity^89–91^, and lower metabolic rate^92^ than during Summer. Because molecular and physiological events associated with hibernation are similar among mammals^36,41,44,93^, the within active season variation in epigenetic aging rate may occur in other mammals. For instance, free-living arctic ground squirrels begin dropping body temperature 45 days before hibernation^94^, 13-lined ground squirrels drops food consumption by 55% prior to hibernation^95^, and some species exhibit short and shallow torpor bouts before and after hibernation^96^.

DNA methylation (DNAm) aging in marmots was related to genes involved in several developmental and differentiation processes—as seen in other mammals^17,19,72,97^. This common enrichment across mammals implies an evolutionary conservation in the biological processes underpinning aging. This inference has been further reinforced by a recent study developing ECs capable of accurately predicting chronological age in distantly related species and, in theory, in any mammal species^70^. These “universal clocks” for eutherians can be used in any tissue sample and are developed from CpG sites located in conserved genomic regions across mammals^53^.

Seasonally dynamic methylation levels were identified in 47 CpG sites. Although few CpGs were identified in our analysis per site, the effect of season was detected by the EPM algorithm, which represents methylation changes in all sites correlated (*r* > 0.7) with chronological age^63,64^. Thus, seasonality probably influences many more CpGs in common with aging than we were able to detect. Nevertheless, many of the top season-related sites were proximal to genes with circannual patterns in other species. For instance, AUTS2 is differently expressed among seasons and within hibernation in brown adipose tissue of 13-lined ground squirrels^98^ and its proximal CpGs are differentially methylated in blood and liver throughout the reproductive season of great tits^99^. JARID2 is differentially expressed within hibernation in the cerebral cortex of 13-lined ground squirrels^100^ and seasonally expressed in human peripheral blood mononuclear cells^101^. RUFY3 is differentially expressed between active and hyperphagia phases in the subcutaneous adipose tissue of grizzly bears^102^ and is close to season-related CpGs in great tits^103^. Methylation levels of sites close to FILIP1, AHDC1, ARHGEF12, ZNF521, CTNNA1 and AUTS2 vary seasonally in great tits^103^. ARHGEF12 is also upregulated in songbirds exhibiting migratory behavior^104^. The expression of these genes may thus be of some importance to species with seasonal behavior, including in hibernating and non-hibernating species.

Since hibernation depends on the synchrony of all regulatory stages^45^ and profoundly alters physiology, most pathways are affected by season in hibernating species. However, little is known about the molecular regulation of seasonal rhythms, and our results imply a role for DNA methylation in regulating some circannual processes, as previously suggested^105^. Seasonal changes in central carbon metabolism and immune responses are expected because immune function is downregulated during hibernation^106^, and the reliance on carbohydrates as energy source is switched for lipid metabolism^44,45^. Remarkably, the circadian clock system was enriched by CpGs related to seasonality. Seasonal changes in photoperiod are encoded in the circadian clock, and modify gene expression in core-clock genes as well as in clock-controlled genes^107–109^.

In sum, our main finding was the little to no change in epigenetic aging during hibernation. While hibernation may increase longevity by protecting individuals from predators and diseases^37^, we suggest that the biological processes involved in hibernation are important contributors to the long lifespan seen in hibernators. Since the reduction of metabolic rates is reached through similar molecular and biochemical patterns across the animal kingdom^110^, the wide inter- and intra-specific variation of torpor use in nature should be explored for more insights about the interplay between aging and torpor.

## Data availability

Epigenetic data will be deposited in Gene Expression Omnibus.

## Author contributions

GMP, SH and DTB conceived the study. GMP, JGAM, CF and AH analyzed data. GMP, AH, DTB, JGAM and SH wrote the manuscript. The remaining authors helped with data generation, statistical analysis and critical feedback. All authors reviewed and edited the article.

## Acknowledgments

We are grateful for all the marmoteers who diligently collected the data over the years, and for the Blumstein, Horvath and Wayne labs for supportive feedback. Thanks also for the helpful insights from the UCLA Statistical Consulting Group, in special to Andy Lin and Siavash Jalal.

## References

1. Flatt, T. A new definition of aging? Front. Genet. 3, 148 (2012).

2. Berdasco, M. & Esteller, M. Hot topics in epigenetic mechanisms of aging: 2011. Aging Cell 11, 181–186 (2012).

3. Jylhävä, J., Pedersen, N. L. & Hägg, S. Biological Age Predictors. EBioMedicine 21, 29–36 (2017).

4. Wagner, K. H., Cameron-Smith, D., Wessner, B. & Franzke, B. Biomarkers of aging: From function to molecular biology. Nutrients 8, 338 (2016).

5. Field, A. E. et al. DNA Methylation Clocks in Aging: Categories, Causes, and Consequences. Mol. Cell 71, 882–895 (2018).

6. Horvath, S. et al. Decreased epigenetic age of PBMCs from Italian semi-supercentenarians and their offspring. Aging 7, 1159–1170 (2015).

7. Nussey, D. H., Froy, H., Lemaitre, J. F., Gaillard, J. M. & Austad, S. N. Senescence in natural populations of animals: Widespread evidence and its implications for bio-gerontology. Ageing Res. Rev. 12, 214–225 (2013).

8. Johnson, T. E. Recent results: Biomarkers of aging. Exp. Gerontol. 41, 1243–1246 (2006).

9. Horvath, S. DNA methylation age of human tissues and cell types. Genome Biol. 14, R115 (2013).

10. Hannum, G. et al. Genome-wide Methylation Profiles Reveal Quantitative Views of Human Aging Rates. Mol Cell 49, 359–367 (2013).

11. Unnikrishnan, A. et al. The role of DNA methylation in epigenetics of aging. Pharmacol. Ther. 195, 172–185 (2019).

12. Bocklandt, S. et al. Epigenetic Predictor of Age. PLoS One 6, e14821 (2011).

13. Horvath, S. & Raj, K. DNA methylation-based biomarkers and the epigenetic clock theory of ageing. Nat. Rev. Genet. 19, 371–384 (2018).

14. Polanowski, A. M., Robbins, J., Chandler, D. & Jarman, S. N. Epigenetic estimation of age in humpback whales. Mol. Ecol. Resour. 14, 976–987 (2014).

15. Petkovich, D. A. et al. Using DNA Methylation Profiling to Evaluate Biological Age and Longevity Interventions. Cell Metab. 25, 954–960 (2017).

16. Stubbs, T. M. et al. Multi-tissue DNA methylation age predictor in mouse. Genome Biol. 18, 68 (2017).

17. Wang, T. et al. Epigenetic aging signatures in mice livers are slowed by dwarfism, calorie restriction and rapamycin treatment. Genome Biol. 18, 57 (2017).

18. Ito, G., Yoshimura, K. & Momoi, Y. Analysis of DNA methylation of potential age-related methylation sites in canine peripheral blood leukocytes. J. Vet. Med. Sci. 79, 745–750 (2017).

19. Thompson, M. J., von Holdt, B., Horvath, S. & Pellegrini, M. An epigenetic aging clock for dogs and wolves. Aging 9, 1055–1068 (2017).

20. Lowe, R. et al. Ageing-associated DNA methylation dynamics are a molecular readout of lifespan variation among mammalian species. Genome Biol. 19, 22 (2018).

21. Zannas, A. S. et al. Lifetime stress accelerates epigenetic aging in an urban, African American cohort: Relevance of glucocorticoid signaling. Genome Biol. 16, 266 (2015).

22. Zaghlool, S. B. et al. Association of DNA methylation with age, gender, and smoking in an Arab population. Clin. Epigenetics 7, 6 (2015).

23. Gao, X., Zhang, Y., Breitling, L. P. & Brenner, H. Relationship of tobacco smoking and smoking-related DNA methylation with epigenetic age acceleration. Oncotarget 7, 46878–46889 (2016).

24. Marioni, R. E. et al. The epigenetic clock and telomere length are independently associated with chronological age and mortality. Int. J. Epidemiol. 45, 424–432 (2016).

25. Marioni, R. E. et al. DNA methylation age of blood predicts all-cause mortality in later life. Genome Biol. 16, 25 (2015).

26. Perna, L. et al. Epigenetic age acceleration predicts cancer, cardiovascular, and all-cause mortality in a German case cohort. Clin. Epigenetics 8, 64 (2016).

27. Chen, B. H. et al. DNA methylation-based measures of biological age: meta-analysis predicting time to death. Aging 8, 1844–1859 (2016).

28. Christiansen, L. et al. DNA methylation age is associated with mortality in a longitudinal Danish twin study. Aging Cell 15, 149–154 (2016).

29. Horvath, S. & Levine, A. J. HIV-1 infection accelerates age according to the epigenetic clock. J. Infect. Dis. 212, 1563–1573 (2015).

30. Horvath, S. et al. Accelerated epigenetic aging in Down syndrome. Aging Cell 14, 491–495 (2015).

31. Parrott, B. B. & Bertucci, E. M. Epigenetic Aging Clocks in Ecology and Evolution. Trends Ecol. Evol. 34, 767–770 (2019).

32. Wagner, W. Epigenetic aging clocks in mice and men. Genome Biol. 18, 107 (2017).

33. Wang, T. et al. Quantitative Translation of Dog-to-Human Aging by Conserved Remodeling of the DNA Methylome. Cell Syst. 11, 1–10 (2020).

34. Wilkinson, G. S. & Adams, D. M. Recurrent evolution of extreme longevity in bats. Biol. Lett. 15, 20180860 (2019).

35. Austad, S. N. Comparative biology of aging. J. Gerontol. A Biol. Sci. Med. Sci. 64, 199–201 (2009).

36. Wu, C. W. & Storey, K. B. Life in the cold: Links between mammalian hibernation and longevity. BioMol. Concepts 7, 41–52 (2016).

37. Turbill, C., Bieber, C. & Ruf, T. Hibernation is associated with increased survival and the evolution of slow life histories among mammals. Proc. R. Soc. B 278, 3355–3363 (2011).

38. Chen, Y. et al. Mechanisms for increased levels of phosphorylation of elongation factor-2 during hibernation in ground squirrels. Biochemistry 40, 11565–11570 (2001).

39. Knight, J. E. et al. mRNA stability and polysome loss in hibernating Arctic ground squirrels (*Spermophilus parryii*). Mol. Cell. Biol. 20, 6374–6379 (2000).

40. Yan, J., Barnes, B. M., Kohl, F. & Marr, T. G. Modulation of gene expression in hibernating arctic ground squirrels. Physiol. Genomics 32, 170–181 (2008).

41. Van Breukelen, F. & Martin, S. L. Molecular adaptations in mammalian hibernators: unique adaptations or generalized responses? J. Appl. Physiol. 92, 2640–2647 (2002).

42. Morin, P. & Storey, K. B. Evidence for a reduced transcriptional state during hibernation in ground squirrels. Cryobiology 53, 310–318 (2006).

43. van Breukelen, F. & Martin, S. L. Reversible depression of transcription during hibernation. J. Comp. Physiol. B Biochem. Syst. Environ. Physiol. 172, 355–361 (2002).

44. Carey, H. V., Andrews, M. T. & Martin, S. L. Mammalian hibernation: Cellular and molecular responses to depressed metabolism and low temperature. Physiol. Rev. 83, 1153–1181 (2003).

45. Al-attar, R. & Storey, K. B. Suspended in time: Molecular responses to hibernation also promote longevity. Exp. Gerontol. 134, 110889 (2020).

46. Azzu, V. & Valencak, T. G. Energy Metabolism and Ageing in the Mouse: A Mini-Review. Gerontology 63, 327–336 (2017).

47. Schrack, J. A., Knuth, N. D., Simonsick, E. M. & Ferrucci, L. ‘IDEAL’ aging is associated with lower resting metabolic rate: The baltimore longitudinal study of aging. J. Am. Geriatr. Soc. 62, 667–672 (2014).

48. Armitage, K. B., Blumstein, D. T. & Woods, B. C. Energetics of hibernating yellow-bellied marmots (*Marmota flaviventris*). Comp. Biochem. Physiol. - A Mol. Integr. Physiol. 134, 101–114 (2003).

49. Armitage, K. B. Phylogeny and patterns of energy conservation in marmots. in Molecules to migration: the pressures of life (eds. Morris, S. & Vosloo, A.) 591–602 (Bologna: Medimond Publishing, 2008).

50. Blumstein, D. T. Yellow-bellied marmots: insights from an emergent view of sociality. Philos. Trans. R. Soc. B 368, 20120349 (2013).

51. Armitage, K. B. Reproductive strategies of yellow-bellied marmots: energy conservation and differences between the sexes. J. Mammal. 79, 385–393 (1998).

52. Armitage, K. B. & Downhower, J. F. Demography of yellow-bellied marmot populations. Ecology 55, 1233–1245 (1974).

53. Arneson, A. et al. A mammalian methylation array for profiling methylation levels at conserved sequences. bioRxiv doi: 10.1101/2021.01.07.425637 (2021) doi:https://doi.org/10.1101/2021.01.07.425637.

54. Zhou, W., Triche, T. J., Laird, P. W. & Shen, H. SeSAMe: reducing artifactual detection of DNA methylation by Infinium BeadChips in genomic deletions. Nucleic Acids Res. 46, e123 (2018).

55. Snir, S., VonHoldt, B. M. & Pellegrini, M. A Statistical Framework to Identify Deviation from Time Linearity in Epigenetic Aging. PLoS Comput. Biol. 12, e1005183 (2016).

56. Snir, S., Wolf, Y. I. & Koonin, E. V. Universal Pacemaker of Genome Evolution. PLoS Comput. Biol. 8, e1002785 (2012).

57. Snir, S., Farrell, C. & Pellegrini, M. Human epigenetic ageing is logarithmic with time across the entire lifespan. Epigenetics 14, 912–926 (2019).

58. Farrell, C., Snir, S. & Pellegrini, M. The Epigenetic Pacemaker: modeling epigenetic states under an evolutionary framework. Bioinformatics 36, 4662–4663 (2020).

59. Zou, H. & Hastie, T. Regularization and variable selection via the elastic net. J. R. Stat. Soc. B 67, 301–320 (2005).

60. Friedman, J., Hastie, T. & Tibshirani, R. Regularization Paths for Generalized Linear Models via Coordinate Descent. J. Stat. Softw. 33, 1–22 (2010).

61. Snir, S. & Pellegrini, M. An epigenetic pacemaker is detected via a fast conditional expectation maximization algorithm. Epigenomics 10, 695–706 (2018).

62. Wood, S. N. Fast stable restricted maximum likelihood and marginal likelihood estimation of semiparametric generalized linear models. J. R. Statist. Soc. B. 73, 3–36 (2011).

63. R Core Team. R: A language and environment for statistical computing. R Foundation for Statistical Computing (2020).

64. RStudio Team. RStudio: Integrated Development Environment for R. RStudio, Inc. (2019).

65. Van Rossum, G. & Drake, F. L. Python 3 Reference Manual. (CreateSpace, 2009).

66. Kluyver, T. et al. Jupyter Notebooks—a publishing format for reproducible computational workflows. Positioning and Power in Academic Publishing: Players, Agents and Agendas (IOS Press, 2016). doi:10.3233/978-1-61499-649-1-87.

67. Wickham, H. ggplot2: Elegant Graphics for Data Analysis. (Springer-Verlag, 2016).

68. Kassambara, A. ggpubr: ‘ggplot2’ Based Publication Ready Plots. https://cran.r-project.org/package=ggpubr (2020).

69. Mclean, C. Y. et al. GREAT improves functional interpretation of cis-regulatory regions. Nat Biotechnol 28, 495–501 (2010).

70. Mammalian Consortium et al. Universal DNA methylation age across mammalian tissues. bioRxiv doi: 10.1101/2021.01.18.426733 (2021).

71. Benjamini, Y. & Hochberg, Y. Controlling the False Discovery Rate: A Practical and Powerful Approach to Multiple Testing. J. R. Statist. Soc. B 57, 289–300 (1995).

72. De Paoli-Iseppi, R. et al. Measuring animal age with DNA methylation: From humans to wild animals. Front. Genet. 8, 106 (2017).

73. Armitage, K. B. Reproductive competition in female yellow-bellied marmots. in Adaptive strategies and diversity in marmots (eds. Ramousse, R., Allainé, D. & Le Berre, M.) 133–142 (International Marmot Network, 2003).

74. Marioni, R. E. et al. Tracking the epigenetic clock across the human life course: A meta-analysis of longitudinal cohort data. J. Gerontol. A Biol. Sci. Med. Sci. 74, 57–61 (2019).

75. El Khoury, L. Y. et al. Systematic underestimation of the epigenetic clock and age acceleration in older subjects. Genome Biol. 20, 283 (2019).

76. Kilgore, D. L. & Armitage, K. B. Energetics of Yellow-Bellied Marmot Populations. Ecology 59, 78–88 (1978).

77. Armitage, K. B. Social and population dynamics of yellow-bellied marmots: results from longterm research. Annu. Rev. Ecol. Syst. 22, 379–407 (1991).

78. Webb, D. R. Environmental harshness, heat stress, and *Marmota flaviventris*. Oecologia 44, 390–395 (1980).

79. Armitage, K. B. Evolution of sociality in marmots. J. Mammal. 80, 1–10 (1999).

80. Allainé, D. Sociality, mating system and reproductive skew in marmots: Evidence and hypotheses. Behav. Processes 51, 21–34 (2000).

81. Arnold, W. The evolution of marmot sociality: II. Costs and benefits of joint hibernation. Behav. Ecol. Sociobiol. 27, 239–246 (1990).

82. Keil, G., Cummings, E. & Magalhães, J. P. Being cool: how body temperature influences ageing and longevity. Biogerontology 16, 383–397 (2015).

83. Means, L. W., Higgins, J. L. & Fernandez, T. J. Mid-life onset of dietary restriction extends life and prolongs cognitive functioning. Physiol. Behav. 54, 503–508 (1993).

84. Speakman, J. R. & Mitchell, S. E. Caloric restriction. Mol. Aspects Med. 32, 159–221 (2011).

85. Walford, R. L. & Spindler, S. R. The response to calorie restriction in mammals shows features also common to hibernation: A cross-adaptation hypothesis. Journals Gerontol. Biol. Sci. 52A, B179–B183 (1997).

86. Conti, B. et al. Transgenic mice with a reduced core body temperature have an increased life span. Science 314, 825–828 (2006).

87. Conti, B. Considerations on temperature, longevity and aging. Cell. Mol. Life Sci. 65, 1626–1630 (2008).

88. Gribble, K. E., Moran, B. M., Jones, S., Corey, E. L. & Mark Welch, D. B. Congeneric variability in lifespan extension and onset of senescence suggest active regulation of aging in response to low temperature. Exp. Gerontol. 114, 99–106 (2018).

89. Johns, D. W. & Armitage, K. B. Behavioral ecology of alpine yellow-bellied marmots. Behav. Ecol. Sociobiol. 5, 133–157 (1979).

90. Armitage, K. B. Social behaviour of a colony of the yellow-bellied marmot (*Marmota flaviventris*). Anim. Behav. 10, 319–331 (1962).

91. Armitage, K. B. Vernal behaviour of the yellow-bellied marmot (*Marmota flaviventris*). Anim. Behav. 13, 59–68 (1965).

92. Armitage, K. B., Melcher, J. C. & Ward, J. M. Oxygen consumption and body temperature in yellow-bellied marmot populations from montane-mesic and lowland-xeric environments. J. Comp. Physiol. B 160, 491–502 (1990).

93. Villanueva-Cañas, J. L., Faherty, S. L., Yoder, A. D. & Albà, M. M. Comparative genomics of mammalian hibernators using gene networks. Integr. Comp. Biol. 54, 452–462 (2014).

94. Sheriff, M. J., Williams, C. T., Kenagy, G. J., Buck, C. L. & Barnes, B. M. Thermoregulatory changes anticipate hibernation onset by 45 days: data from free-living arctic ground squirrels. J. Comp. Physiol. B 182, 841–847 (2012).

95. Schwartz, C., Hampton, M. & Andrews, M. T. Hypothalamic gene expression underlying pre-hibernation satiety. Genes, Brain Behav. 14, 310–318 (2015).

96. Geiser, F. Metabolic rate and body temperature reduction during hibernation and daily torpor. Annu. Rev. Physiol. 66, 239–274 (2004).

97. Maegawa, S. et al. Widespread and tissue specific age-related DNA methylation changes in mice. Genome Res. 20, 332–340 (2010).

98. Hampton, M., Melvin, R. G. & Andrews, M. T. Transcriptomic analysis of brown adipose tissue across the physiological extremes of natural hibernation. PLoS One 8, e85157 (2013).

99. Lindner, M. et al. Temporal changes in DNA methylation and RNA expression in a small song bird: within- and between-tissue comparisons. BMC Genomics 22, 36 (2021).

100. Schwartz, C., Hampton, M. & Andrews, M. T. Seasonal and Regional Differences in Gene Expression in the Brain of a Hibernating Mammal. PLoS One 8, e58427 (2013).

101. Dopico, X. C. et al. Widespread seasonal gene expression reveals annual differences in human immunity and physiology. Nat. Commun. 6, 7000 (2015).

102. Jansen, H. T. et al. Hibernation induces widespread transcriptional remodeling in metabolic tissues of the grizzly bear. Commun. Biol. 2, 336 (2019).

103. Viitaniemi, H. M. et al. Seasonal Variation in Genome-Wide DNA Methylation Patterns and the Onset of Seasonal Timing of Reproduction in Great Tits. Genome Biol. Evol. 11, 970–983 (2019).

104. Johnston, R. A., Paxton, K. L., Moore, F. R., Wayne, R. K. & Smith, T. B. Seasonal gene expression in a migratory songbird. Mol. Ecol. 25, 5680–5691 (2016).

105. Stevenson, T. J. Epigenetic Regulation of Biological Rhythms: An Evolutionary Ancient Molecular Timer. Trends Genet. 34, 90–100 (2018).

106. Bouma, H. R., Carey, H. V. & Kroese, F. G. M. Hibernation: the immune system at rest? J. Leukoc. Biol. 88, 619–624 (2010).

107. Coomans, C. P., Ramkisoensing, A. & Meijer, J. H. The suprachiasmatic nuclei as a seasonal clock. Front. Neuroendocrinol. 37, 29–42 (2015).

108. Sumová, A., Bendová, Z., Sládek, M., Kováčikova, Z. & Illnerová, H. Seasonal Molecular Timekeeping Within the Rat Circadian Clock. Physiol. Res. 53, S167–S176 (2004).

109. Meijer, J. H., Michel, S. & Vansteensel, M. J. Processing of daily and seasonal light information in the mammalian circadian clock. Gen. Comp. Endocrinol. 152, 159–164 (2007).

110. Storey, K. B. & Storey, J. M. Metabolic rate depression in animals: Transcriptional and translational controls. Biol. Rev. 79, 207–233 (2004).

